# Soybean Growth and Nodulation Under Mars-Like Magnesium Sulfate Concentrations

**DOI:** 10.64898/2026.09.10.750807

**Authors:** Caio Felipe de Miranda-Costa, Ana Paula Avelino, Carlos Takeshi Hotta

**Author notes:** **Correspondence:** Carlos Takeshi Hotta.

## Abstract

Colonising Mars requires sustainable food production systems that use local resources. However, the Martian regolith presents significant challenges for plant cultivation. We investigated whether soybean (*Glycine max* (L.) Merr.) could grow and form symbiotic nodules with *Bradyrhizobium japonicum* in MGS-1, a high-fidelity Martian regolith simulant. Soybean plants grown in an MGS-1:vermiculite mix (1:2) and supplemented with a nutrient solution were shorter, produced fewer leaves, and accumulated less biomass than sand-grown plants. More importantly, no nodulation occurred in MGS-1, despite successful nodulation in sand-grown controls. To identify factors inhibiting nodulation, we tested increasing concentrations of MgSO₄ (50–150 mM), a major salt in MGS-1, and osmotic stress (300 mM Mannitol). While both treatments reduced nodule numbers, nodulation persisted across all MgSO₄ concentrations and under osmotic stress, indicating that neither factor fully accounts for the absence of nodules in MGS-1. The shoot-to-root dry weight ratio increased under MgSO₄ stress but decreased under mannitol, suggesting distinct stress responses. Our findings demonstrate that soybean can survive in MGS-1 with nutrient supplementation and an additional substrate to improve MGS-1 compactness and porosity. Still, soybean grown in MGS-1 exhibits severe developmental constraints and complete inhibition of biological nitrogen fixation. Overcoming these limitations will require strategies to mitigate salinity, alkalinity, and physical compaction, as well as engineering symbiotic conditions compatible with sustainable agriculture on Mars.

## Introduction

Colonising Mars will require continuous delivery of food to astronauts, which is very costly, unless food can be grown on the Red Planet (Creech, Guidi & Elburn 2022; Noble *et al*. 2024). In an ideal scenario, plants could sustain the survival of a crew far from Earth, producing food as part of their edible bioweight and simultaneously contributing to various ecological services, such as air purification and water recycling (Fu *et al*. 2016), in addition to promoting the psychological well-being of space explorers (Odeh & Guy 2017). However, although plants have been grown in space for more than 50 years, no successful food production system has yet been developed to sustain astronauts (Zabel, Bamsey, Schubert & Tajmar 2016).

Mars is the fourth planet in the Solar System. Its atmosphere is composed mainly of carbon dioxide (CO₂, ∼95%), nitrogen (N₂, ∼3%) and argon (∼Ar, ∼2%), plus trace gases such as oxygen and carbon monoxide (Mahaffy *et al*. 2013; Starr & Muscatello 2020; Vandaele *et al*. 2024), in addition to having an atmospheric pressure that represents 0.6% of Earth’s atmosphere (Vandaele *et al*. 2024). This thin atmosphere and lack of an ozone layer mean UV radiation is significantly higher on Mars than on Earth (Cockell & Andrady 1999). Mars’ average temperature is around −65 °C, although equatorial regions can reach up to 20 °C during summer days (Piqueux *et al*. 2024). Gravity on Mars corresponds to 0.375 g (Rahaim & Czysz 2008). A Martian day lasts 24 h 37 min, and the planet is 229 million kilometres (1.5 astronomical units) from the Sun, with an axial tilt of 25° compared to Earth’s 23.4°. Such similarities as day length and axial tilt between the two planets mean that Mars not only has a photoperiod similar to that of Earth, but also experiences seasons comparable to those we know, with the difference being the duration of each season, since a Martian year lasts 687 Earth days (Martínez *et al*. 2017).

Mars lacks soil because its surface contains no organic matter or biological activity. It has a layer of loose rock and sediments that covers bedrock, called planetary regolith (Long-Fox, Landsman, Easter, Millwater & Britt 2023). The Martian regolith has a generally basaltic mineralogical profile that is broadly similar across the planet at the bulk elemental level. However, significant local and regional mineralogical heterogeneity exists (Ehlmann & Edwards 2014), with silicon dioxide (SiO₂), iron oxides (FeO/Fe₂O₃), aluminium oxide (Al₂O₃), and magnesium oxide (MgO) as the major components (Yen *et al*. 2005; McSween, Taylor & Wyatt 2009; Meslin *et al*. 2013). Magnesium sulphate (MgSO_4_) is estimated to account for more than 50% by weight in the soluble salt fraction of the regolith (Toner, Catling & Light 2014; Chevrier, Gil-Lozano, Dehouck & Altheide 2024). Perchlorate (ClO₄⁻) concentrations in the saline phase are also noteworthy, as they are toxic to most living beings (Toner *et al*. 2014; Chevrier *et al*. 2024).

Several challenges remain in growing food on Mars. Even with greenhouses and artificial lighting to mitigate low solar radiation, temperature extremes, and low atmospheric pressure, Martian regolith may still be unsuitable for plants. Perchlorates can act as strong oxidisers, posing hazards to greenhouse equipment and human health if dust becomes airborne (Nicholson *et al*. 2012; Davila, Willson, Coates & McKay 2013). Both potassium (K) and bioavailable nitrogen (N) are present at very low levels in the regolith. Phosphorus (P) availability is also limited because high iron content tends to bind P to insoluble forms (Wamelink, Frissel, Krijnen, Verwoert & Goedhart 2014; Wamelink & Pouwels 2024).

Perchlorate levels in the regolith can be reduced by converting it into safer ions, such as chloride (Misra, Smith, Garner & Loureiro 2021). Phosphorus availability can be mitigated by adjusting the regolith pH or using microorganisms to mobilise it from the regolith (Hausrath *et al*. 2024). One way to address low concentrations of N and K is to export them from Earth, but this is expensive. Another solution for N is to use nitrogen-fixing bacteria, either free-living or symbionts (Harris, Dobbs, Atkins, Ippolito & Stewart 2021; Tarasashvili, Elbakidze, Doborjginidze & Gharibashvili 2023).

Although it is currently impossible to use Martian regolith in experiments, artificial substrates that mimic its key features have been developed, such as JSC Mars-1A, MMS-1, MMS-2, and more recently, MGS-1 (Allen *et al*. 1998; Peters *et al*. 2008; Cannon, Britt, Smith, Fritsche & Batcheldor 2019). Researchers used these simulated regoliths to grow several species, such as lettuce (*Lactuca sativa* L.) and potato (*Solanum tuberosum* L.) (Duri et al., 2022). In general, the simulated regoliths lack essential nutrients, especially N, to support plant growth. They also show poor water-holding capacity, which can be mitigated by adding nutrient solutions and mixing the simulated regoliths with compost or other compounds (Wamelink *et al*. 2014; Duri *et al*. 2022; Caporale *et al*. 2024).

We tested whether soybean (*Glycine max* (L.) Merr.) grown in simulated Mars regolith (MGS-1) exhibited developmental changes compared with sand controls and whether it could nodulate. Soybeans are a well-known source of protein, oils, plant-based milk, and even biofuels. It is a diploid legume (2n = 40) important for food production, with a seed protein content of 40% (Singh & Hymowitz 1999; Lee *et al*. 2021). Its life cycle consists of two phases: vegetative and reproductive stages. The vegetative stages are defined by the number of nodes with fully expanded leaves. All V1 to Vn stages have true leaves (leaflet edges no longer touching) that are trifoliolate and produced singularly on different nodes, with these leaves alternating on the stem. The reproductive stages are divided into four parts: flowering, pod development, seed development, and plant and seed maturation (Endres & Carcedo 2025).

Nodules are unique and specialised root structures that result from a symbiotic association between plants in the family Leguminosae and soil bacteria of the genus *Rhizobium* (Zilli, Santa Cruz, Polizio, Tomaro & Balestrasse 2011). These structures capture atmospheric N_2_ and convert it into compounds that are bioavailable to plants. Soybean nodules are colonised by *Bradyrhizobium japonicum* (Kaneko *et al*. 2002). Although air pressure may be an issue on Mars, it may be possible to use the N2 in the Martian atmosphere to generate biologically active N.

In this study, we show that soybean plants grown on a mix of MGS-1:vermiculite (1:2) supplemented with nutrient solution exhibited developmental problems compared with sand controls. In this substrate, individuals are subjected not only to salinity stress but also to compaction, aeration limitations, and substrate pH. Nevertheless, all plants grew in the MGS-1 simulated regolith, demonstrating that it does not fully inhibit growth. However, it does completely inhibit nodulation. We also show that soybean plants grown in sand can tolerate up to 150 mM MgSO₄. Nodulation occurred at all MgSO₄ concentrations in plants grown in sand, indicating that although high salt concentrations affect nodulation, they are not fully inhibitory. Plants also nodulate under osmotic stress, which explains part of the effects observed under high MgSO₄ concentrations, but not all.

## Materials and methods

### Plant material and seed germination

Soybean (*Glycine max* (L.) Merr.) cultivar BRS 284, a high-yielding variety from Embrapa (Embrapa Soja, Paraná, Brazil), was used in all experiments. Seeds were disinfected by immersion in 70% ethanol (v/v) for 3 min, rinsed five times in sterile distilled water, and imbibed in sterile distilled water for 2 h. Seeds were sown on square bioassay polystyrene dishes (243 mm) with filter paper moistened with sterile distilled water, and incubated in a growth chamber (12 h light/12 h dark photoperiod, 65% relative humidity, and 27 °C) for 4 days.

### Treatments and growth conditions

In the first experiment, three-day-old seedlings were transferred to plastic pots containing either Mars Global Simulant 1 (MGS-1) (Space Resource Technologies, Florida, USA) mixed with vermiculite (Brasil Minérios, Goiás, Brazil) or washed medium sand (Sand) (Areião ABC, São Paulo, Brazil) mixed with vermiculite, both at a 1:2 ratio (v:v). MGS-1 is a Martian regolith simulant based on analyses conducted by NASA’s Curiosity rover at Rocknest in Gale Crater on Mars, reproducing current knowledge of the physical and chemical properties of Martian soil, except for perchlorates (Cannon et al., 2019). Sand was used as a substrate control because it is also nutrient-poor, which limits nitrogen bioavailability for soybean plants and thus favours nodulation. Still, it has a larger particle size than MGS-1. We mixed both substrates with vermiculite to increase water retention and reduce substrate compaction. Before seedling transfer, each substrate mixture was moistened with tap water until saturation to create water channels for subsequent sub-irrigation. The pots were distributed into trays (per treatment) and sub-irrigated twice a week, alternating between nutrient solution and tap water (Visscher et al., 2010). The nutrient solution used was Murashige and Skoog (Sigma-Aldrich) half-strength, supplemented with 2-morpholinoethanesulfonic acid monohydrate (Sigma-Aldrich), pH 5.75 (Visscher et al., 2010). In the following experiments, we transferred three-day-old seedlings to plastic pots containing sand mixed with vermiculite. The substrate mixture was prepared and sub-irrigated as previously described, except for the nutrient solution composition. In the second experiment, to investigate the impact of increasing MgSO₄ concentrations on soybean growth and nodulation, the nutrient solution was supplemented with 50, 100, and 150 mM MgSO₄ · 7H₂O (Sigma-Aldrich). In the third experiment, to test whether MgSO₄ effects were due to osmotic stress, the nutrient solution was supplemented with 150 mM MgSO₄ · 7H₂O (Sigma-Aldrich) or 300 mM Mannitol (Sigma-Aldrich). In both experiments, the control plants were sub-irrigated with the nutrient solution without additions. In all experiments, soybean plants were either inoculated or non-inoculated with *Bradyrhizobium japonicum*. All trays were maintained in a growth tent under a 12 h light/12 h dark photoperiod and 25 ± 5 °C until harvest.

### Plant inoculation with Bradyrhizobium japonicum

Plant inoculation was carried out with the liquid inoculant RIZOKOP (SEMIA 5079 and 5080, 7 × 10^9^ CFU/mL) (Koppert, Brazil). The inoculation process consisted of adding the liquid inoculant to the substrate mixture at three time points: immediately after seedling transfer (5 mL), the following week (10 mL), and after the emergence of the first trifoliolate leaf (10 mL). We increased the RIZOKOP volume to ensure the inoculant penetrated the substrate.

### Harvests and plant growth parameters

Plants were harvested at 28 days (first and third experiments) and 36 days (second experiment) after imbibition (DAI). During harvests, we recorded soybean plant morphology by digital imaging, manually counted leaf and nodule numbers, and measured shoot height with a graduated ruler. Soybean plant root systems were gently washed with tap water to remove the substrate mixture and allow nodule counting. After superficially drying the roots, plants were divided into shoots and roots and immediately weighed to determine fresh weight (FW). Samples were oven-dried at 60 °C for 72 h and weighed again to determine dry weight (DW). Water content (WC) was calculated using the following formula: WC = {(FW – DW)/FW} × 100. For biochemical determinations, the two youngest trifoliolate leaves and the upper root portion were frozen and kept at −80 °C until quantification.

### Quantification of total soluble proteins and photosynthetic pigments

Samples were ground in liquid nitrogen, and one hundred milligrams of the pulverised sample were used to determine the total soluble protein (TSP) content. TSPs were extracted from soybean leaves and roots using 1.5 mL of 100 mM Tris-HCl buffer (pH 7.0). Samples were mixed by inversion and then centrifuged at 10,000 g for 20 min at 4 °C. Supernatants were collected, and the pellets were re-extracted twice with 1 mL of the extraction buffer, yielding a total of 3.5 mL. We used the Bradford assay (Bio-Rad) to determine TSP concentration, with bovine serum albumin (Sigma-Aldrich) as the standard (Bradford 1976). Samples were measured at 595 nm using a plate spectrophotometer (BMG Labtech). TSP content was expressed in mg·g^-1^ DW.

Total chlorophylls and carotenoids were extracted from fifty milligrams of pulverised leaves with 7.5 mL of ice-cold 80% acetone (v/v) under reduced luminosity. Samples were protected from light, briefly vortexed, and kept at 4 °C for 72 h. Afterwards, samples were centrifuged at 10,000 g for 10 min at 4 °C, and the supernatants were used to estimate total chlorophyll (Arnon 1949) and carotenoid (Jaspars 1965) contents. Measurements were performed at 450, 645, and 663 nm using a plate spectrophotometer (BMG Labtech). Total chlorophyll and carotenoid concentrations were expressed in mg·g^-1^ FW.

### Experimental design and data analysis

We conducted the experiments in a completely randomised design. The experimental unit consisted of a tray with six soybean plants. We analysed the results using Analysis of Variance (ANOVA) and two-way ANOVA, followed by Tukey tests for multiple pairwise comparisons (p < 0.05) (Supp. Tables 1, 2 and 4). In the first experiment, substrate mixture type (MGS-1 or sand) and inoculation status (inoculated or non-inoculated) were the independent variables, while plant growth parameters were the fixed variables. In the following experiments, MgSO_4_ and mannitol concentrations and inoculation status were the independent variables. To evaluate the effect of MgSO₄ concentration and inoculum type on length, we fitted a multiple linear regression model using ordinary least squares (OLS). The model included MgSO₄ as a continuous fixed effect and Inoculum as a categorical fixed effect (Supp. Table 3). All codes for statistical analysis and figures are available at: https://github.com/LabHotta/Nodulation_Mars.

## Results

### MGS-1 inhibits growth, nodulation, and alters biomass allocation in soybean

To assess the effects of a Martian regolith simulant on soybean development, seedlings of the BRS 284 cultivar were grown on a mix of MGS-1:vermiculite (1:2), with a mix of sand: vermiculite (1:2) used as a substrate control. Because both substrates were nutrient-poor, we applied a nutrient solution once a week and used a *Bradyrhizobium japonicum* liquid inoculant to promote nodulation in half of the seedlings. MGS-1-grown plants were smaller, accumulated less biomass, and were unable to develop nodules compared with sand-grown plants (Figure 1). In fact, plants grown in MGS-1 were 33% shorter (Figure 1C), had 39% fewer trifoliolate leaves (Supp. Figure 1A), and 67% less biomass (Figure 1D, Supp. 1D) than plants grown on sand. Additionally, while inoculated plants developed 13.8 ± 8.7 nodules (n = 6) when grown on sand, no nodules were detected on inoculated MGS-1-grown plants (Figure 1B; H-K). Although inoculated sand-grown plants had nodules in their roots (Figure 1I), inoculation did not affect most analysed growth parameters (Figure 1, Supp. Figure 1 and Supp. Table 1). Curiously, even without visible nodules (Figure 1K), inoculation increased leaf TSP content by 21% (Figure 1L) in MGS-1-grown plants in relation to non-inoculated plants grown on the same substrate.

**Figure 1.**
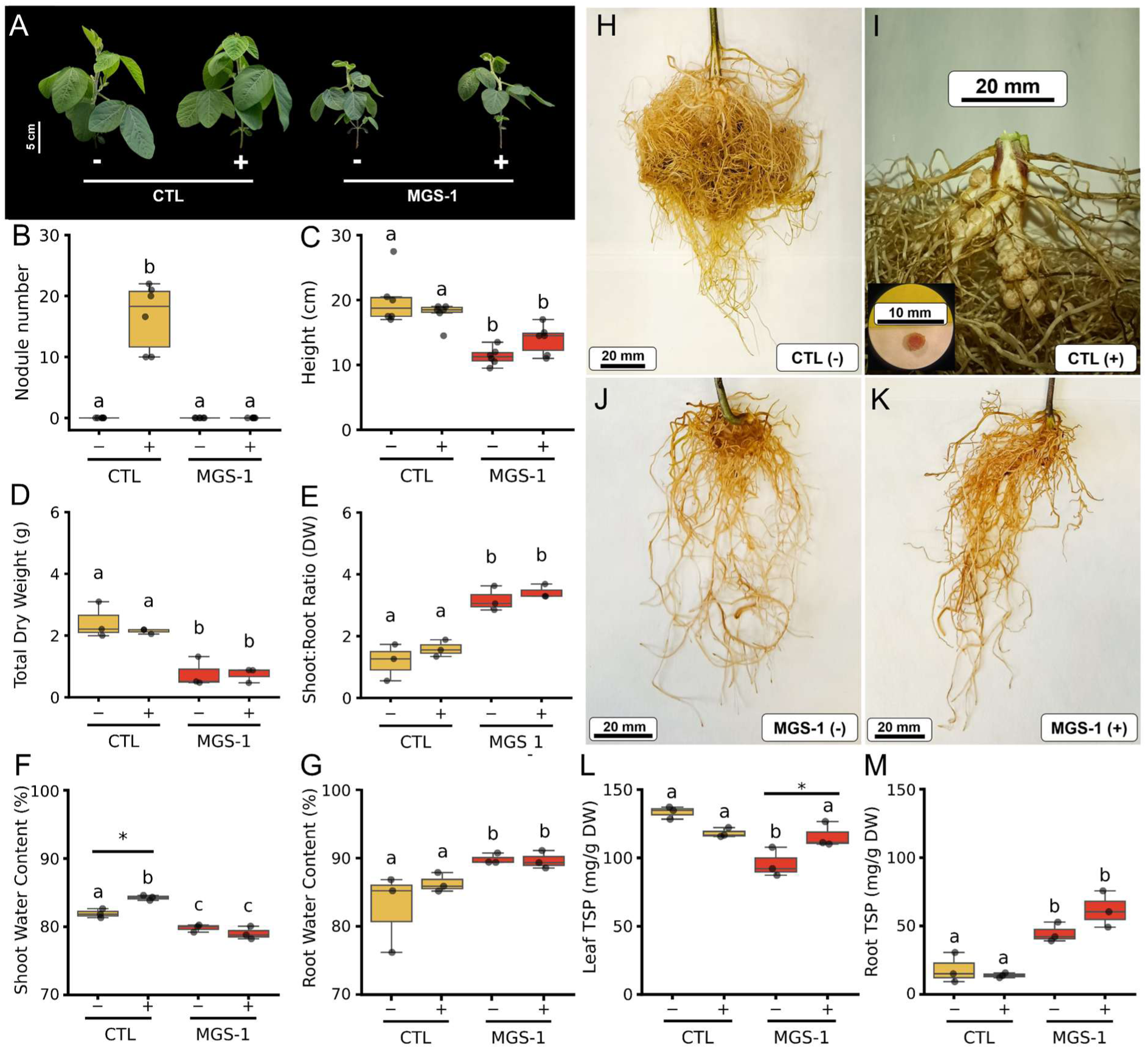
Effects of Mars Global Simulant (MGS-1) on growth, nodulation, and protein content of soybean, with or without *Bradyrhizobium japonicum* inoculation. (**A**) Representative shoot morphology. **(H–K)** Corresponding root systems: **(H)** non-inoculated control (CTL); **(I)** inoculated CTL (inset: nodule cross-section); **(J)** non-inoculated MGS-1; and **(K)** inoculated MGS-1. Quantitative data are shown for **(B)** nodule number, **(C)** plant height, **(D)** total dry weight, **(E)** shoot-to-root dry weight ratio, **(F)** shoot water content, **(G)** root water content, **(L)** leaf total soluble protein (TSP), and **(M)** root TSP. Plants were grown in either a 1:2 sand: vermiculite mixture (CTL, yellow bars) or a 1:2 MGS–1:vermiculite mixture (MGS–1, red bars), with inoculation status indicated by (+) or (–) (n = 6 for panels B–C; n = 3 for panels D–M). Lowercase letters denote significant differences among substrate treatments (two–way ANOVA with Tukey’s post hoc test, p < 0.05). At the same time, asterisks (*) indicate significant differences between inoculated and non–inoculated plants within the same substrate (p < 0.05).

MGS-1 strongly affected soybean root development, altering shoot-to-root biomass allocation (Figure 1E). Plants grown on MGS-1 accumulated 82% less dry weight in roots than plants grown on sand (Supp. Figure 1C). Consequently, a 2.4-fold increase in the shoot-to-root dry weight ratio was observed under the same growth conditions, indicating that these plants increased biomass allocation to shoots (Figure 1E). Differences in water and TSP content allocation were also found between sand- and MGS-1-grown plants (Figure 1F, G, L and M). While plants grown on sand had 4% more water in shoots than plants grown on MGS-1, MGS-1-grown plants had 6% more water in roots in relation to the sand-grown controls (Figure 1F and G). Following a similar pattern, leaves of sand-grown plants exhibited 24% more TSP than leaves of non-inoculated MGS-1-grown plants (Figure 1L). By contrast, in the roots of plants grown on MGS-1, TSP content increased 3.3-fold compared with sand-grown plants (Figure 1M). Despite that, no alterations were seen in total chlorophyll or carotenoid contents (Supp. Figure 1J and K). Collectively, the low performance of soybean plants grown on MGS-1 and the absence of nodulation are indicative that these plants were under stress.

### High MgSO₄ concentrations inhibit growth but do not prevent nodulation

To test whether high Mg-sulfate concentrations in MGS-1 could explain the observed differences (Cannon et al., 2019; Long-Fox et al., 2023), we grew inoculated and uninoculated plants on sand supplemented with 50-150 mM MgSO_4_ (Figures 2, 3 and Supp. 2). Regression analysis revealed a significant relationship between MgSO₄ concentration and most of the tested variables, with most effects being negative (Supp. Table 2). For example, in both inoculated and non-inoculated plants, plant height was 4.3% lower for every 10 mM MgSO_4_, while leaf number was 8.5% lower for every 10 mM MgSO_4_ (Figure 3B and C).

**Figure 2.**
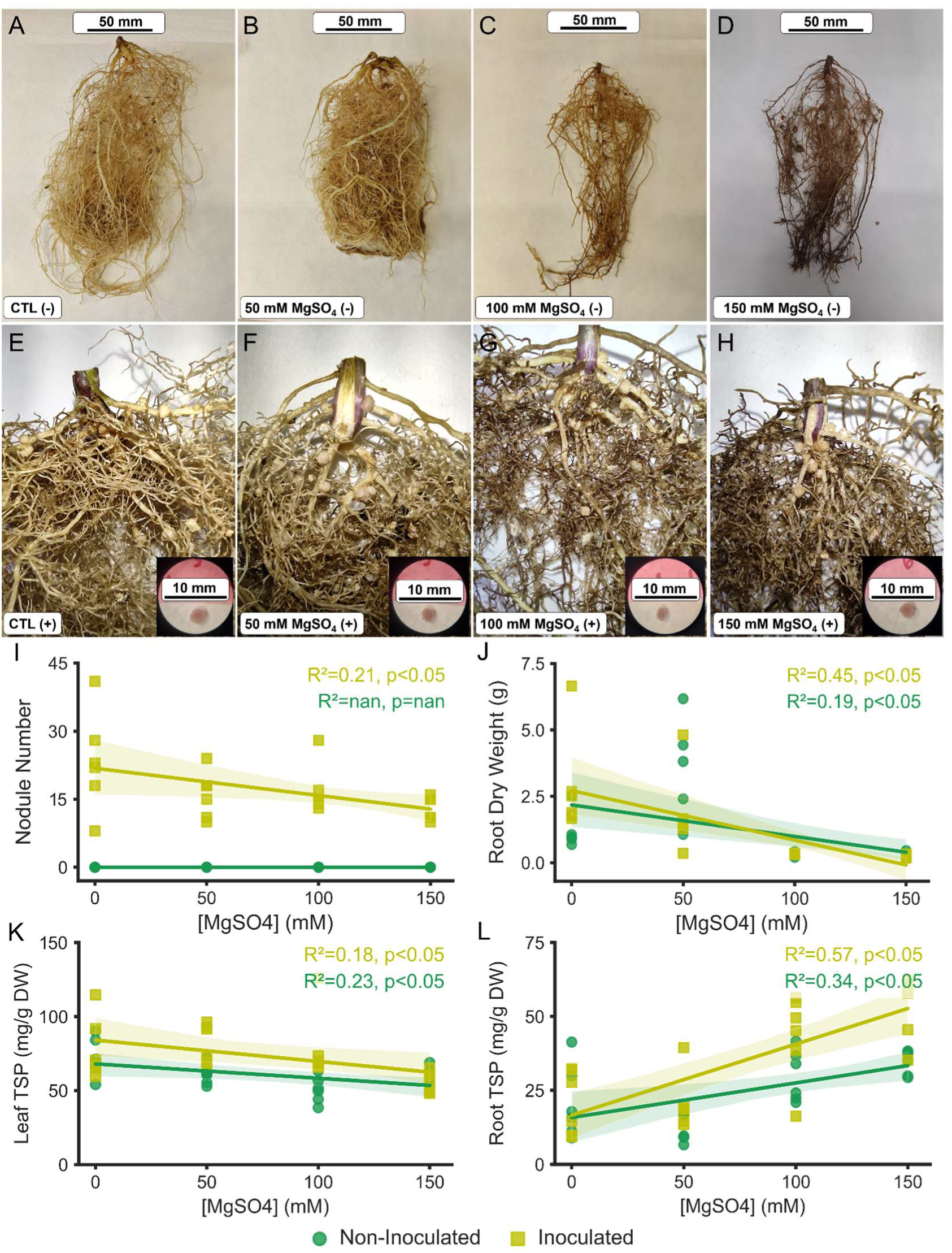
Influence of increasing concentrations of MgSO_4_ on nodule formation in soybean plants, inoculated or not with *Bradyrhizobium japonicum*. (**A–D**) Representative root systems from non-inoculated plants. (**E–H**) Representative root systems from plants inoculated with *Bradyrhizobium japonicum*; the lower-right inset shows a histological cross-section of a nodule excised from the corresponding root. Quantitative data are presented for (**I**) nodule number, (**J**) root dry weight, (**K**) leaf total soluble protein (TSP), and (**L**) root TSP. Solid regression lines (light green, inoculated; dark green, non-inoculated) indicate trends across concentrations (n = 6 biological replicates per condition). R² and p-values (derived from linear regression) are provided in the upper-right of each quantitative panel to indicate fit and statistical significance.

**Figure 3.**
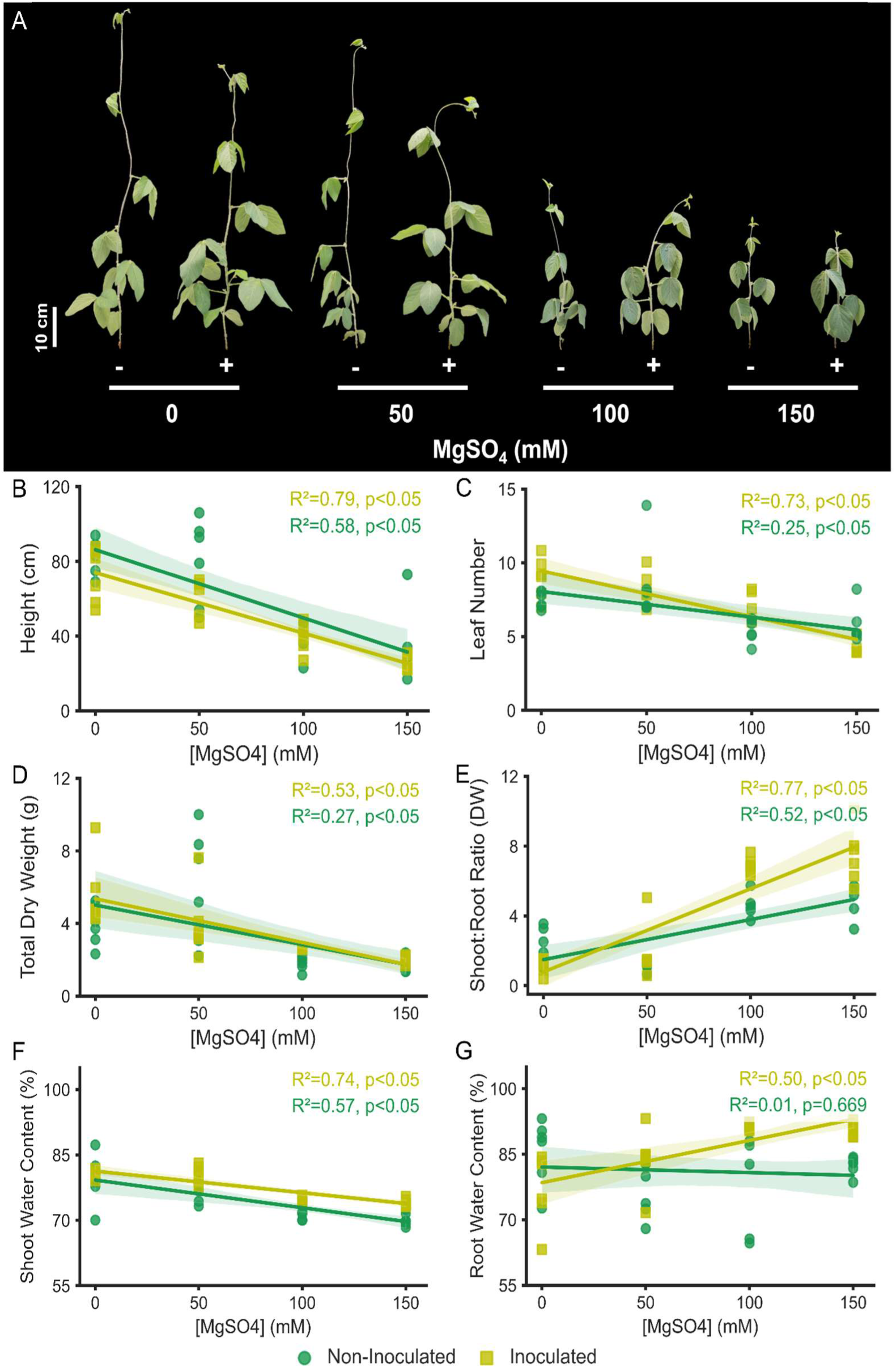
Influence of increasing concentrations of MgSO_4_ on growth parameters in soybean plants, inoculated or not with *Bradyrhizobium japonicum*. **(A)** Representative shoot morphology. Quantitative data are presented for (**B**) plant height, **(C)** leaf number, **(D)** total dry weight, **(E)** shoot-to-root dry weight ratio, **(F)** shoot water content, and **(G)** root water content. Solid regression lines (light green, inoculated; dark green, non-inoculated) indicate trends across concentrations (n = 6 biological replicates per condition). R² and p-values (derived from linear regression) are provided in the upper-right of each quantitative panel to indicate fit and statistical significance.

The number of nodules was negatively affected by MgSO_4_ (Figure 2A-I). While inoculated plants treated with 0 mM MgSO_4_ had 23.3 ± 10.9 nodules (n = 6), plants treated with 150 mM MgSO_4_ had 13.0 ± 2.6 nodules (n = 6). Overall, each 10 mM MgSO4 reduced nodule number by 13.3%. Thus, although MgSO_4_ reduced nodule numbers, it was unlikely to account for the complete absence of nodulation in MGS-1. Cross sections of nodules also show reddish interiors due to leghemoglobin, a colourimetric indicator of nitrogen-fixing activity (Figure 2E-H).

Despite differences in the number of nodules, root dry weight in inoculated and non-inoculated plants was similarly affected by MgSO_4_, with a 94% reduction observed at 150 mM MgSO_4_ (Figure 2J). In contrast, the inoculated plants had higher root TSP than non-inoculated plants when treated with MgSO_4_ (Figure 2L). In fact, root TSP increased at a higher rate in inoculated plants than in non-inoculated plants, increasing by 16.0% per 10 mM MgSO_4_ in inoculated plants, compared with 8.4% per 10 mM MgSO_4_ in non-inoculated plants. In contrast, leaf TSP was similarly reduced in inoculated and non-inoculated plants (Figure 2K).

The total dry weight of inoculated and non-inoculated plants is equally affected, with a 66% reduction in plants treated with MgSO_4_ (Figure 3D). In contrast, the shoot: root dry weight ratio increases with MgSO_4_ concentration, as the root dry weight is more affected by MgSO_4_ than the shoot dry weight (Figures 2J, 3E, Supp. Figure 2A). In non-inoculated plants, the shoot: root ratio of dry weight increases from 2.31 ± 0.99 (n = 6) in 0 mM MgSO_4_ to 4.94 ± 0.96 (n = 6) in 150 mM MgSO_4_. In inoculated plants, the shoot: root dry weight ratio goes from 1.19 ± 0.47 (n = 6) in 0 mM MgSO_4_ to 7.47 ± 1.62 (n = 6) in 150 mM MgSO_4_. The shoot water content decreases by 0.72% for every 10 mM increase in MgSO_4_ (Figure 3F), but the root water content remains unaffected in non-inoculated plants (Figure 3G). However, in inoculated plants, root water content increases by 0.51% per 10 mM increase in MgSO_4_.

Principal component analysis of data from the MGS-1 and MgSO4 experiments showed clear separation between stressed and non-stressed plants (Figures 4A and Supp. Figure 3). Among the non-stressed plants, the control plants in each experiment and plants treated with 50 mM MgSO4 were identified, confirming that this MgSO4 concentration was not stressful to the plants. Among the stressed plants (treated with MGS-1, 100 mM MgSO_4,_ or 150 mM MgSO_4_), plants with nodules were separated from those without, suggesting that nodulation affected plants’ responses to stress (Figure 4A). We generated a correlation matrix using growth parameters from nodulated plants (Figure 4B). Nodule number positively correlated with leaf number, shoot dry weight and root dry weight, and negatively with root TSP.

**Figure 4.**
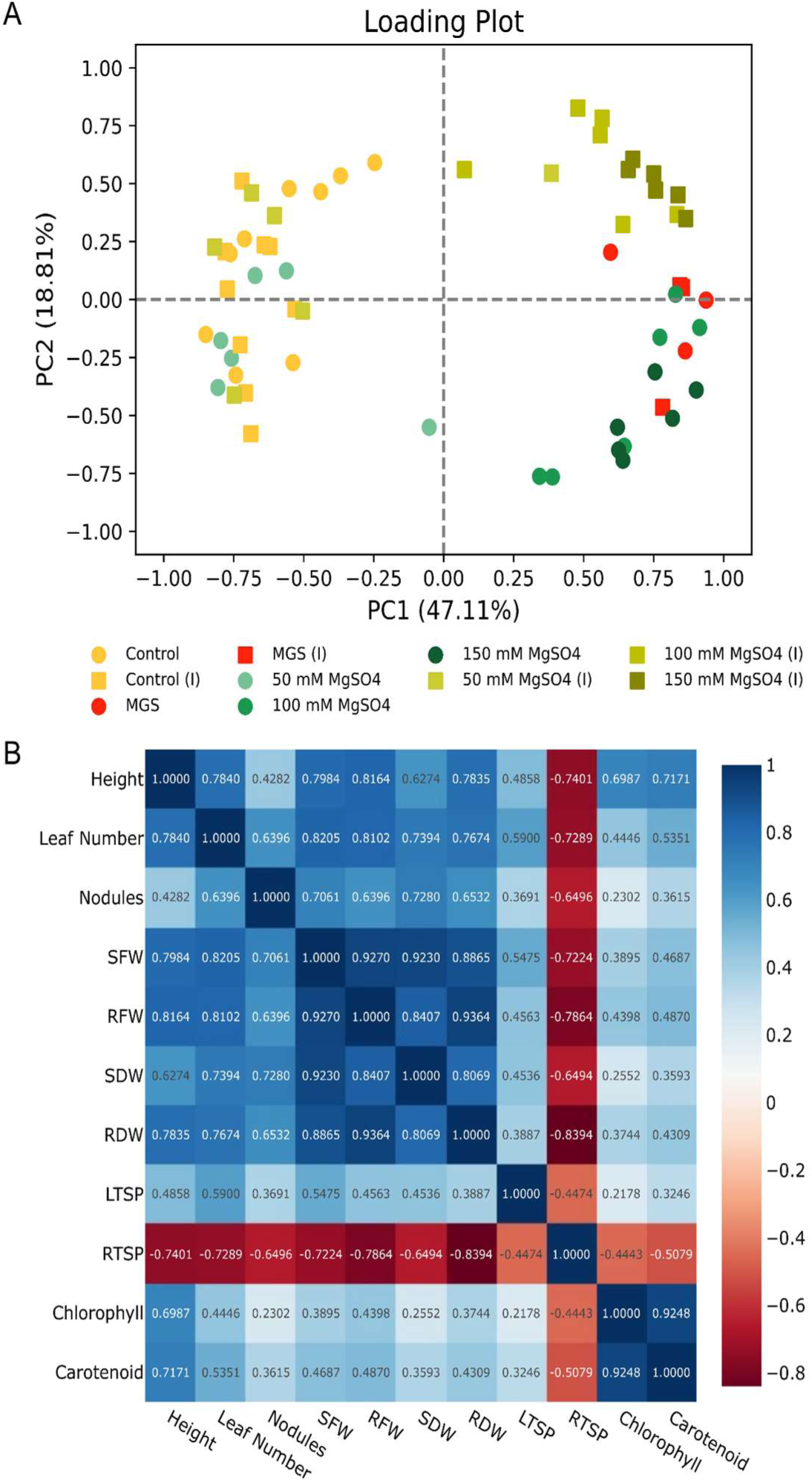
Principal component analysis (PCA) and correlation matrix of growth and physiological parameters in soybean. **(A)** The biplot displays the distribution of treatments across the first two principal components (PC1 = 58.67% and PC2 = 18.81% of total variance). Variables included in the analysis were nodule number, plant height, total dry weight, shoot-to-root dry weight ratio, shoot and root water content, and total soluble protein (TSP) in leaves and roots. Treatments are colour-coded by substrate type: control (yellow), MgSO₄-amended (green gradient), and Mars Global Simulant (MGS-1; red). Inoculation status with *Bradyrhizobium japonicum* is indicated by squares (inoculated) or circles (non-inoculated) symbols. Each data point represents a biological replicate. **(B)** Correlation matrix showing Spearman correlation coefficients between variables in plants that showed nodulation, with deeper colour indicating stronger correlations. Blue shades denote positive correlations, while red shades show negative correlations. Variables include: Height, Leaf Number, Nodules, Shoot Dry Weight (SDW), Root Dry Weight (RDW), Shoot: Root Dry Weight Ratio (DWratio), Leaf Total Soluble Protein (LTSP), Root Total Soluble Protein (RTSP), Chlorophyll, and Carotenoid content.

### Mannitol does not elicit the same responses as MgSO₄ in soybean

The responses observed in the MgSO_4_ experiment could reflect osmotic stress rather than a specific stress response to these ions. To test that, we treated plants with 150 mM MgSO_4_ or 300 mM mannitol (Figures 5A and Supp. Figures 4-6). Both MgSO_4_ and Mannitol decreased the number of nodules in inoculated plants (Figure 5B). While the control plants had 37.0 ± 11.0 nodules (n = 6), plants treated with MgSO_4_ had 24.8 ± 7.8 nodules (n = 6), and plants treated with 300 mM Mannitol had 18.2 ± 7.7 nodules (n = 6). Mannitol was also unable to completely prevent nodule formation in MGS-1 (Supp. Figure 4G).

**Figure 5.**
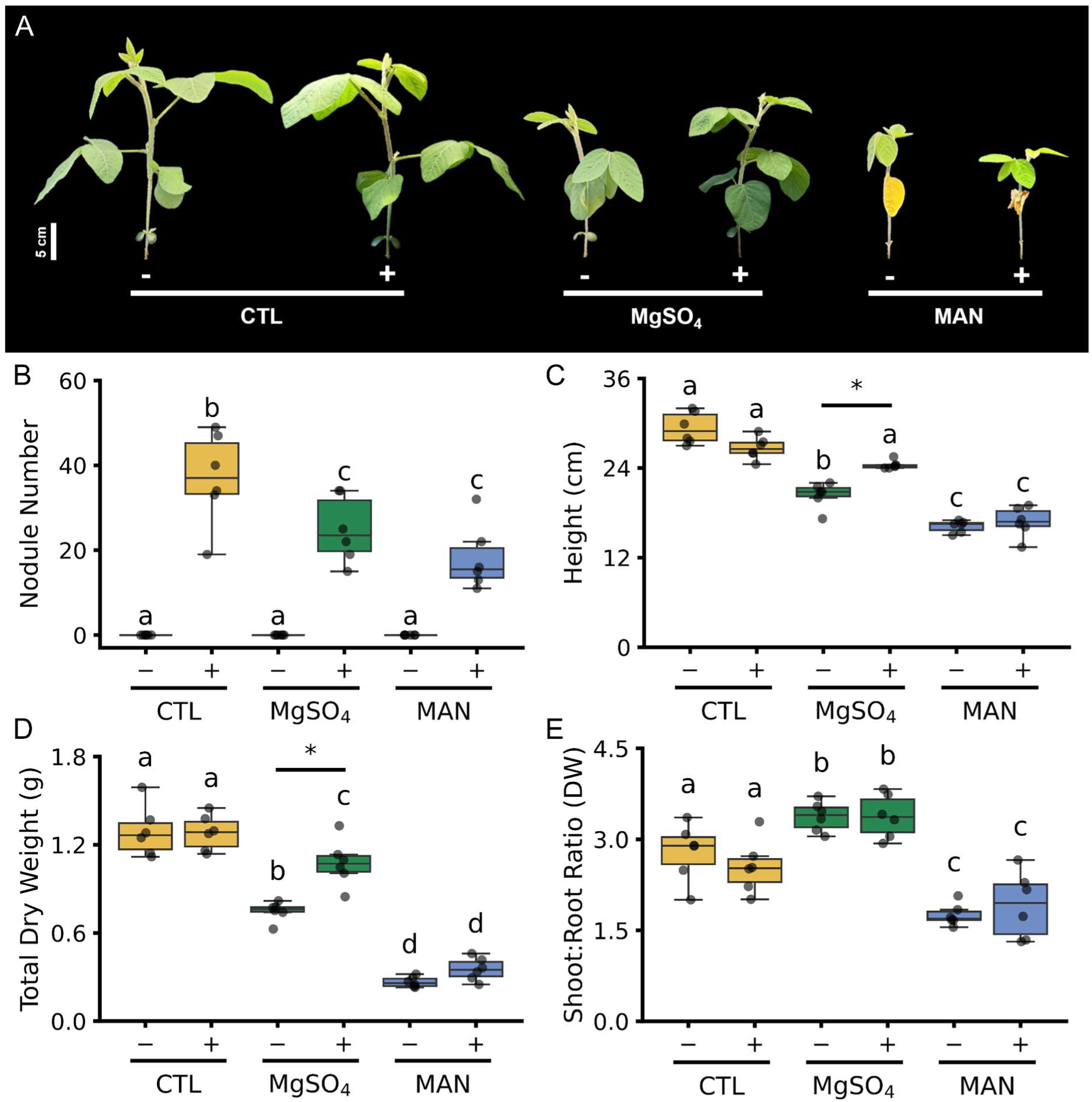
Effects of MgSO_4_ and Mannitol on soybean growth and nodulation, with or without *Bradyrhizobium japonicum* inoculation. (**A**) Representative shoot morphology. Quantitative data show **(B)** nodule number, **(C)** plant height, **(D)** total dry weight, and **(E)** shoot-to-root dry weight ratio. Plants were treated with water (yellow), 150 mM MgSO_4_ (green) and 300 mM Mannitol (blue), with inoculation status indicated by (+) or (–) (n = 6). Lowercase letters denote significant differences among treatments (two–way ANOVA with Tukey’s post hoc test, p < 0.05). At the same time, asterisks (*) indicate significant differences between inoculated and non–inoculated plants within the same substrate (p < 0.05).

Height and total dry height were also decreased in treated plants, but inoculated plants were less affected by 150 mM MgSO_4_ (Figure 5C and D). Non-inoculated plants treated with 150 mM MgSO_4_ were 30.6% shorter and 42.1% lighter than control plants, whilst inoculated plants treated with 150 mM MgSO_4_ were not statistically different in height and were 15.9% lighter. In contrast, plants treated with 300 mM Mannitol were 44.9% shorter and 79.4% lighter than control plants. No difference was found between inoculated and non-inoculated plants (Supp. Table 4).

Most soybean responses to 150 mM MgSO_4_ and 300 mM Mannitol were in the same direction. Both 150 mM MgSO_4_ and 300 mM Mannitol reduced water content in the shoots and roots (Supp. Figure 5C and D). The exception was the shoot: root dry weight ratio (Figure 5E). As in the previous experiment, 150 mM MgSO_4_ increased the shoot-to-root dry weight ratio by 21.1%. However, 300 mM Mannitol reduced it by 37.1% (Figure 5E), suggesting that osmotic stress cannot account for all the effects observed in the 150 mM MgSO_4_ treatment.

## Discussion

Crewed space missions to Mars directly imply long-duration missions. However, Earth-supplied resources become a limiting factor not only because of cost but also because of the time and difficulty inherent to Martian launches (Llorente, Williams & Goold 2018; Duri *et al*. 2022). Therefore, a bioregenerative food system is essential for the success of a long-term mission, especially if *in situ* resources are used (Nelson, Dempster & Allen 2008; Perchonok, Cooper & Catauro 2012; De Micco *et al*. 2023). Thus, using Mars regolith as a substrate for growing plants is essential to the viability of a human presence on the red planet.

Plant growth on Mars regolith using regolith simulants. First experiments using simulated Mars regolith (JSC Mars-1A, Mojave Mars Simulant-1, and MMS-2) showed that plants could grow on these substrates, especially with a source of nutrient supplementation (Wamelink *et al*. 2014; Duri *et al*. 2020; Eichler *et al*. 2021; Wamelink & Pouwels 2024; Gonçalves, Wamelink, Putten & Evers 2024). However, plants grow poorly on Mars Global Simulant-1 (Eichler *et al*. 2021; Chinnannan *et al*. 2024), a new generation of simulated regolith that represents the current understanding of Martian regolith composition. MGS-1 is particularly hydrophobic, so it requires an additional component to improve water retention. (Cannon *et al*. 2019; Bacior *et al*. 2025). When grown directly on MGS-1, both *Arabidopsis thaliana* (L.) Heynh, *Ipomoea batatas* (L.) Lam. (sweet potato), and *Lactuca sativa* L. (lettuce) perished after a few days, even with nutrient supplementation (Eichler *et al*. 2021; Chinnannan *et al*. 2024). When MGS-1 was mixed with another substrate and given external nutrients, survivability increased (Chinnannan *et al*. 2024; Bacior *et al*. 2025; Servetto *et al*. 2026).

We tested whether *Glycine max* (L.) Merr. (soybean) grew on MGS-1 mixed with vermiculite (1:2, v:v) and supplemented with a nutrient solution. We chose soybean for this study because of its protein content and oil production (Li *et al*. 2026). Plants grown in MGS-1 were shorter and accumulated less biomass, but the leaves did not show signs of leaf damage, and chlorophyll content was similar to plants grown on sand (Figure 1A, C and D; Supp. Figures 1D, J and 7). Sweet potato showed a similar biomass reduction at 50% MGS-1 exposure and a 25% decrease in chlorophyll content (Chinnannan *et al*. 2024). Soybean grown in MGS-1 showed an increase in the shoot: root ratio of dry weight (Figure 1E), contrasting with soybean grown in MMS-1, which showed no changes in dry matter partitioning (Caporale *et al*. 2022). Soybean grown in MGS-1 also has lower total soluble protein (TSP) content in the leaves than in the roots (Figure 1L and M). This indicates that, under stress, sink organs maintain essential proteins but show reduced metabolism compared with the source organ, as shown by a 30% reduction in TSP in this tissue. In contrast, roots of plants grown in MGS-1 showed up to a four-fold increase in TSP content (Figure 1M) compared to the sand control. Since this tissue is in direct contact with multiple stressors and therefore must activate metabolic pathways to mitigate toxic effects, such as proline synthesis to prevent water loss and the production of dehydrin family proteins, which protect root cells from reactive oxygen species generated by stress (Khedr, Abbas, Wahid, Quick & Abogadallah 2003; Balasubramaniam, Shen, Esmaeili & Zhang 2023).

Soybeans can form nodules containing *Bradyrhizobium japonicum* that can fix atmospheric nitrogen. Nitrogen fixation by diazotrophic bacteria contributes to the sustainability of long-duration missions by reducing dependence on synthetic nitrogen fertilisers transported from Earth, as these microorganisms convert atmospheric nitrogen into biologically available forms. Consequently, forming effective plant-bacteria symbioses is important for both legume adaptation and the bioremediation of extraterrestrial regolith (Atkin, Pierson, Gentry & Oliveira Santos 2026). Plants grown in MGS-1 inoculated with *Bradyrhizobium japonicum* did not develop nodules (Figure 1B and K). The regolith simulant contains multiple stressors, including high salinity, alkaline pH (>8.0), the absence of rhizosphere microorganisms, and smaller particle size, which makes it more compact (Gonçalves *et al*. 2024; Duri *et al*. 2025). MGS-1 has high concentrations of MgSO_4_ (Cannon et al., 2019; Long-Fox et al., 2023), so we tested if this salt could explain the absence of nodulation in MGS-1. MgSO_4_ inhibited nodule formation, but nodules were still present up to 150 mM MgSO_4_ (Figure 2F-I). We also observed reduced nodulation capacity in soybean plants at 100 mM NaCl, although these are different salts (Nitawaki, Kitabayashi, Mason, Yamamoto & Saeki 2021). We also tested a solution with high osmotic potential (300 mM Mannitol), which did not eliminate nodule formation, although it inhibited nodulation more than 150 mM MgSO_4_ (Figures 5 and Supp. Figure 4). Thus, neither MgSO_4_ nor osmotic stress could fully explain the absence of nodules in MGS-1.

Nodulation was previously successful in *Melilotus officinalis* (L.) Lam (clover) inoculated with *Sinorhizobium meliloti* in JSC-1 (Harris *et al*. 2021). However, nodule formation was reduced by> 75%. Only 1 in 10 *Lathyrus oleraceus* Lam. (= *Pisum sativum* L., pea) inoculated with *Rhizobium leguminosarum* showed nodule formation in MMS-1 (Gonçalves *et al*. 2024). In contrast, the plant model legume *Medicago truncatula* Gaertn. inoculated with *Sinorhizobium meliloti* and *Sinorhizobium medicae* in MMS-1 and MMS-2, developed the same number of nodules as on sand (Rainwater & Mukherjee 2021). Many factors may contribute to poor nodulation on Mars simulant regoliths: pH, excess of salts, osmotic stress and substrate compactness. In our experiments, MGS-1 had pH = 7.8 (Supp. Table 5). Although some *Bradyrhizobium* strains can tolerate pH = 8.4 (Miladinović *et al*. 2024), in this study, the strains used were *B. japonicum* SEMIA 5079 and 5080, for which the recommended pH for the culture medium is between 6.8 and 7.0 (Döbereiner, Andrade & Baldani 1999; Nakei, Venkataramana & Ndakidemi 2022). Thus, the absence of nodulation may result from the high alkalinity of MGS-1 (Gonçalves *et al*. 2024). We tested whether high MgSO₄ concentrations or osmotic stress could explain the absence of nodules in soybean grown in MGS-1 (Figures 2I and 5B). In both treatments, nodule numbers were smaller than in the control, but still present. Thus, these factors don’t fully explain the absence of nodules in plants grown in MGS-1. To mitigate the effects of soil compaction, we attempted to reduce the impact of the MGS-1’s smaller particle size by mixing it with vermiculite, but this was insufficient to promote nodulation. One possibility is inoculating soybeans with Arbuscular Mycorrhizal Fungi (AMF), as this has been reported to bioremediate Lunar Regolith Simulant for another legume species, Cicer arietinum L. (chickpea), from growth to seed (Atkin *et al*. 2026). The compactness of wet MGS-1 also traps ethylene gas. This plant hormone can inhibit Nod factor signal transduction, which is key to establishing symbiosis in *Medicago* for nodulation (Oldroyd, Engstrom & Long 2001).

Nodulation had little impact on the growth parameters of non-stressed plants in our experiments, likely because we used nutrient solutions. Stressed plants incubated with *Bradyrhizobium* had higher total soluble protein in the roots, a higher shoot: root ratio (DW), and higher root water content (Figures 1E, G and M; 2L; 3E and G). Stressed plants that were incubated with *Bradyrhizobium* were grouped separately from stressed plants that were not incubated in a PCA (Figure 4A). Nodulation protection against salt, cold, and drought stress is well documented and may be linked to enhanced antioxidant systems and improved stress-response modulation (Wang *et al*. 2016, 2026; Chakraborty *et al*. 2021).

Salinity stress is an important environmental factor that restricts plant growth and development, particularly by affecting underground organs (Chourasia *et al*. 2022). Supplementation of the soybean cultivation substrate with different concentrations of MgSO₄ altered growth, with decreased plant performance observed at 100 mM MgSO₄ and above (Figures 2 and 3). High soil salt concentrations negatively affect plants by causing osmotic stress, ionic imbalance, toxicity, and reduced photosynthesis (Chourasia *et al*. 2022). Under osmotic stress, plants may exhibit drought-related symptoms because water moves less efficiently from a medium with a lower solute concentration (soil) to one with a higher solute concentration (root cells). This occurs because the salt concentration in the substrate hinders water entry into root cells, leading to stunted growth, wilting, and even plant death (Bhattacharya 2021). Ionic imbalance occurs when excess ions are present outside the cell, which can dehydrate the root and impair processes such as root growth (Chourasia *et al*. 2022).

High concentrations of MgSO₄ were observed to reduce the length, fresh weight, and dry weight of soybean plants (Figures 3B, D and Supp. Figure 2E). Excess magnesium ions have been studied in *Solanum lycopersicum* L. (tomato) plants, and concentrations of 3 mM MgSO₄ have been shown to impair plant growth and the uptake of K⁺ and Ca²⁺ (Li, Liu, Zhang, Gao & Chen 2024). In comparison, the present study used MgSO₄ concentrations up to 50 times higher than those previously reported. Thus, growth parameters were expected to decrease; however, soybean plants grew well up to 50 mM MgSO₄. This may occur because Mg²⁺ plays a central role in key processes of plant development. Magnesium is an essential plant nutrient and is absorbed in amounts similar to those of phosphorus, with a critical content of 4 g/kg in soybeans. Mg²⁺ is mobile within plants and has essential functions such as: (1) production of the chlorophyll molecule; (2) being a component of ribosomes, directly associated with protein synthesis; (3) activation of enzymes such as glutathione synthetase, methionine activator, pyruvate kinase, pyrophosphatase, pyruvate dehydrogenase, ribulose-2P carboxylase, among others; (4) stabilisation of nucleic acid structures; and (5) acting as an essential element for ATP formation (Tian *et al*. 2021). Thus, it can be proposed that a concentration of 50 mM MgSO₄ is high enough for Mg²⁺ to fulfil its functions within plants, yet low enough that natural defence mechanisms, such as the production of proline, soluble sugars, glycine betaine, and polyols (Abd El-Mageed *et al*. 2022), are sufficient to mitigate stress effects. Soybean plant phenology is associated with the number of nodes (Fehr & Caviness 1977). The number of nodes correlates with the physiological parameter of leaf number. Thus, with MgSO₄ supplementation, ontogenetic development was altered in the sand substrate, with the rate halved at 150 mM MgSO₄. This result raises the hypothesis that MgSO₄ stress on soybean phenology is intensified by greater nutritional restriction in the substrate. A common symptom of salinity stress in plants is leaf yellowing (chlorosis). This symptom was not observed in plants supplemented with MgSO₄ (Figures 2A, 5A and Supp. 7), which may be attributable to magnesium’s central role in chlorophyll synthesis (Tian *et al*. 2021). Arabidopsis growth was severely affected by 100 mM MgSO₄. Still, effects could be mitigated by mutation in *CAX1* (*VACUOLAR CATION/PROTON EXCHANGER 1*), a vacuolar H^+^/Ca^2+^ transporter (Cheng, Pittman, Barkla, Shigaki & Hirschi 2003; Visscher *et al*. 2010). Thus, mutations in these transporters could favour plant growth in Mars regolith.

Osmotic stress and high MgSO4 concentrations had similar effects on soybean (Figure 5). However, shoot: root (DW) increased in plants grown on MGS-1 and at high MgSO_4_ concentrations but decreased under 300 mM Mannitol. This suggests that changes in C allocation are a specific effect of MgSO_4_, likely due to Mg²⁺ effects on plant growth and photosynthesis (Tian *et al*. 2021).

Fixing biological N on Mars has other challenges not tested in this work. Mars has a lower percentage of nitrogen in its atmosphere than Earth, which may affect the bacteria’s ability to fix atmospheric nitrogen (Rainwater & Mukherjee 2021). Another obstacle present in Martian regolith but absent in MGS-1 is perchlorate, which is known to be extremely toxic to plant growth (Oze *et al*. 2021). In addition to studies on plant tolerance to these perchlorates, another possible strategy is to use microorganisms for bioremediation, enabling the removal of these ultratoxic components (Misra *et al*. 2021).

Plant growth in the new-generation simulated Mars regolith, MGS-1, is more challenging than in JSC-1, MMS-1, and MMS-2. However, it can be accomplished by adding a nutrient solution and a substrate that improves regolith porosity and bulk compactness, creating spaces that allow water and air to flow. Osmotic stress and MgSO₄ play a central role in plant stress in simulated regolith, as we observed decreases across all parameters analysed, consistent with observations from the MGS-1 experiment; thus, further investigation is needed to isolate additional MGS-1 stressors and to understand how each affects plant growth. Biological nitrogen fixation is a desired trait for a bioregenerative food system. However, nodulation was not observed in MGS-1, even though it was seen in both high MgSO_4_ concentrations and osmotic stress conditions. Several strategies could promote nodulation, including pH adjustment, incorporating larger particles to improve substrate structure, adding organic material, and applying ethylene inhibitors. Another strategy to adapt Mars regolith for agricultural use is to reuse this substrate generation after generation, which may close the gap between an inorganic substrate and proper soil. Improving bioregenerative life support systems requires local resources, such as Mars regolith, and future research should focus on modifying this substrate to make it more amenable to plants and on breeding or engineering plants with increased tolerance to the conditions it presents.

## Acknowledgements

This work received financial support from the São Paulo Research Foundation (FAPESP) under grant no. 22/13970-7 as part of the Program on Global Climate Change. CTH thanks the National Council for Scientific and Technological Development (CNPq) for the Research Productivity Grant (Bolsa de Produtividade em Pesquisa) (309537/2022-3). CFMC received a scholarship from the Conselho Nacional para o Desenvolvimento Científico e Tecnológico (CNPq).

## Author Contributions

Conceptualisation, CFMC and CTH; Methodology, CFC, APA, and CTH; Software, CFMC, APA, and CTH; Formal Analysis, CFMC, APA, and CTH; Investigation, CFMC and APA; Resources, CTH; Data Curation, CFMC and CTH; Writing – Original Draft Preparation, CFMC and CTH; Writing – Review & Editing, CFMC, APA, and CTH; Visualisation, CFMC, APA, and CTH; Supervision, CTH; Project Administration, CTH; Funding Acquisition, CTH.

## Conflict of Interest and Other Ethics Statements

The authors declare no conflicts of interest.

