## Supplemental Figures and Table for "Soybean Growth and Nodulation Under Mars-Like Magnesium Sulfate Concentrations"

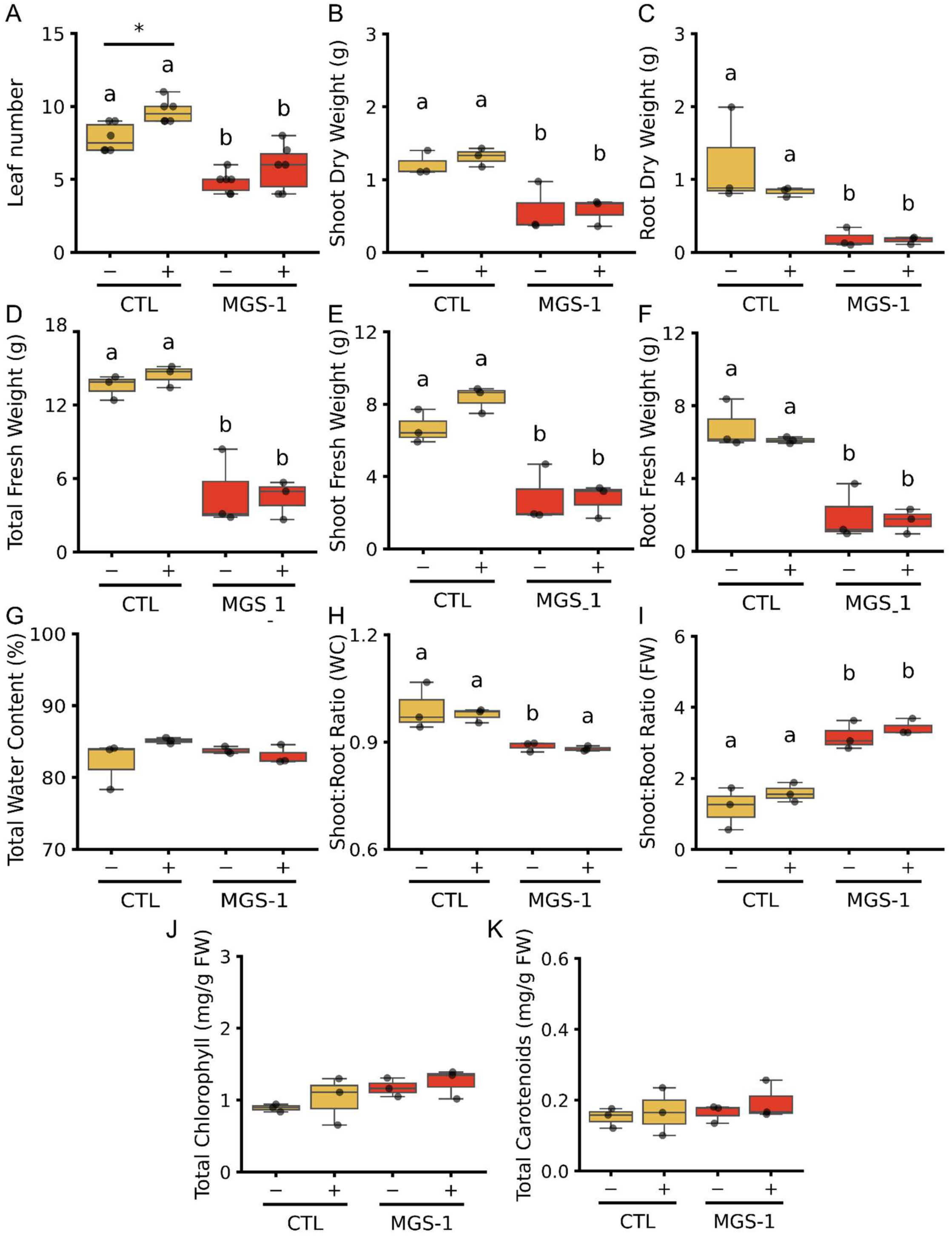


**Supplementary Figure 1. Effects of Mars Global Simulant (MGS-1) on growth parameters and photosynthetic pigment content of soybean plants, inoculated or not with *Bradyrhizobium japonicum*.** **(A)** Leaf number, **(B)** shoot dry weight, **(C)** root dry weight, **(D)** total fresh weight, **(E)** shoot fresh weight, **(F)** root fresh weight, **(G)** total water content, **(H)** shoot-to-root water content ratio, **(I)** shoot-to-root fresh weight ratio, **(J)** total chlorophylls, and **(K)** total carotenoids. Plants were grown in either a 1:2 sand: vermiculite mixture (CTL, yellow bars) or a 1:2 MGS-1: vermiculite mixture (MGS1, red bars), with inoculation status indicated by (+) or (–) (n = 6 for panel A; n = 3 for panels B–K). Lowercase letters denote significant differences among substrate treatments (two-way ANOVA with Tukey's post hoc test, $p < 0.05$). At the same time, asterisks (*) indicate significant differences between inoculated and non-inoculated plants within the same substrate ($p < 0.05$).

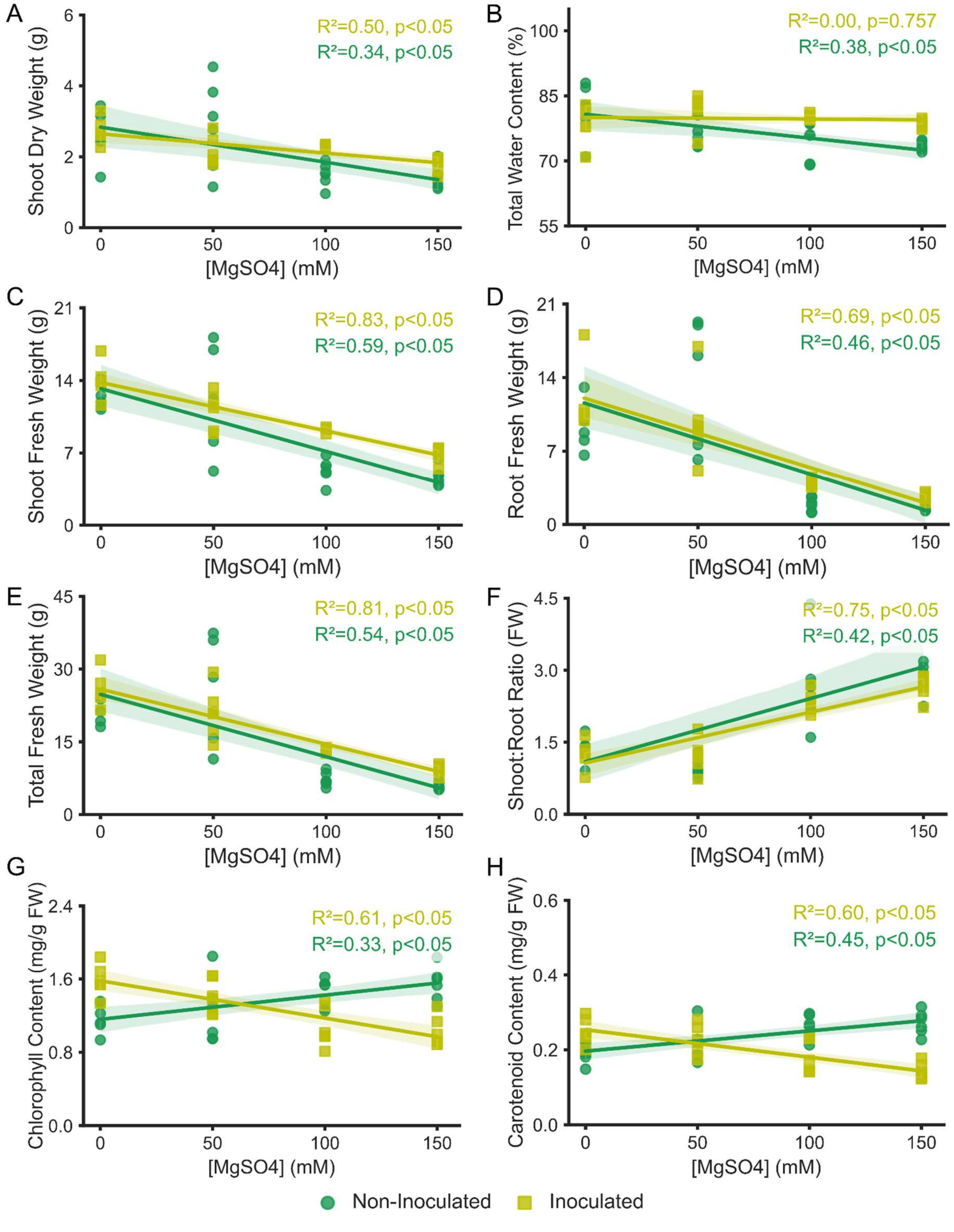


**Supplementary Figure 2. Influence of increasing concentrations of $MgSO_4$ on growth parameters and photosynthetic pigment content of soybean plants, inoculated or not with *Bradyrhizobium japonicum*.** **(A)** Shoot dry weight, **(B)** total water content, **(C)** shoot fresh weight, **(D)** root fresh weight, **(E)** total fresh weight, **(F)** shoot-to-root fresh weight ratio, **(G)** total chlorophylls, and **(H)** total carotenoids. Solid regression lines (light green, inoculated; dark green, non-inoculated) indicate trends across concentrations (n = 6 biological replicates per condition). $R^2$ and p-values (derived from linear regression) are provided in the upper-right of each quantitative panel to indicate fit and statistical significance.

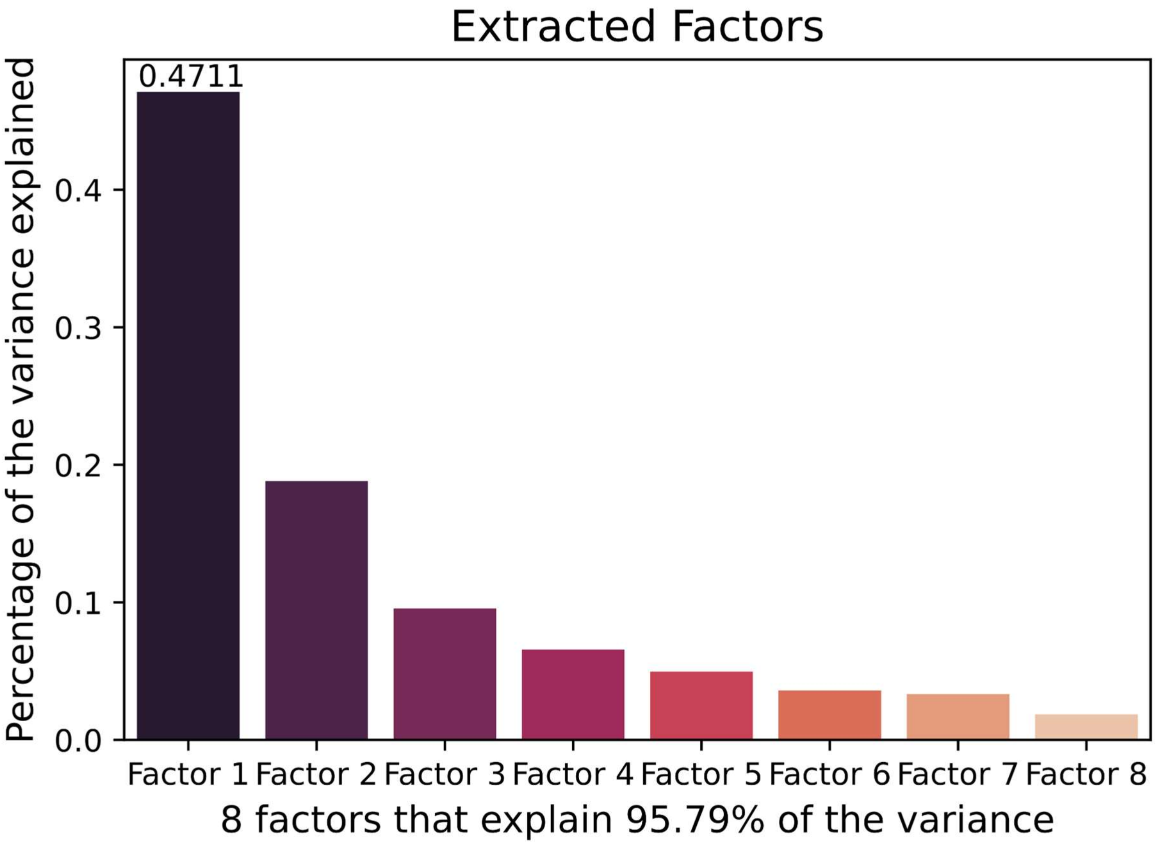


**Supplementary Figure 3 - Factor extraction results.** The bar chart displays the variance explained by each of the 8 factors retained from the analysis. The cumulative percentage (95.79%) is highlighted, confirming that these factors capture most of the information in the original variables, leaving only 4.21% of the variance unaccounted for.

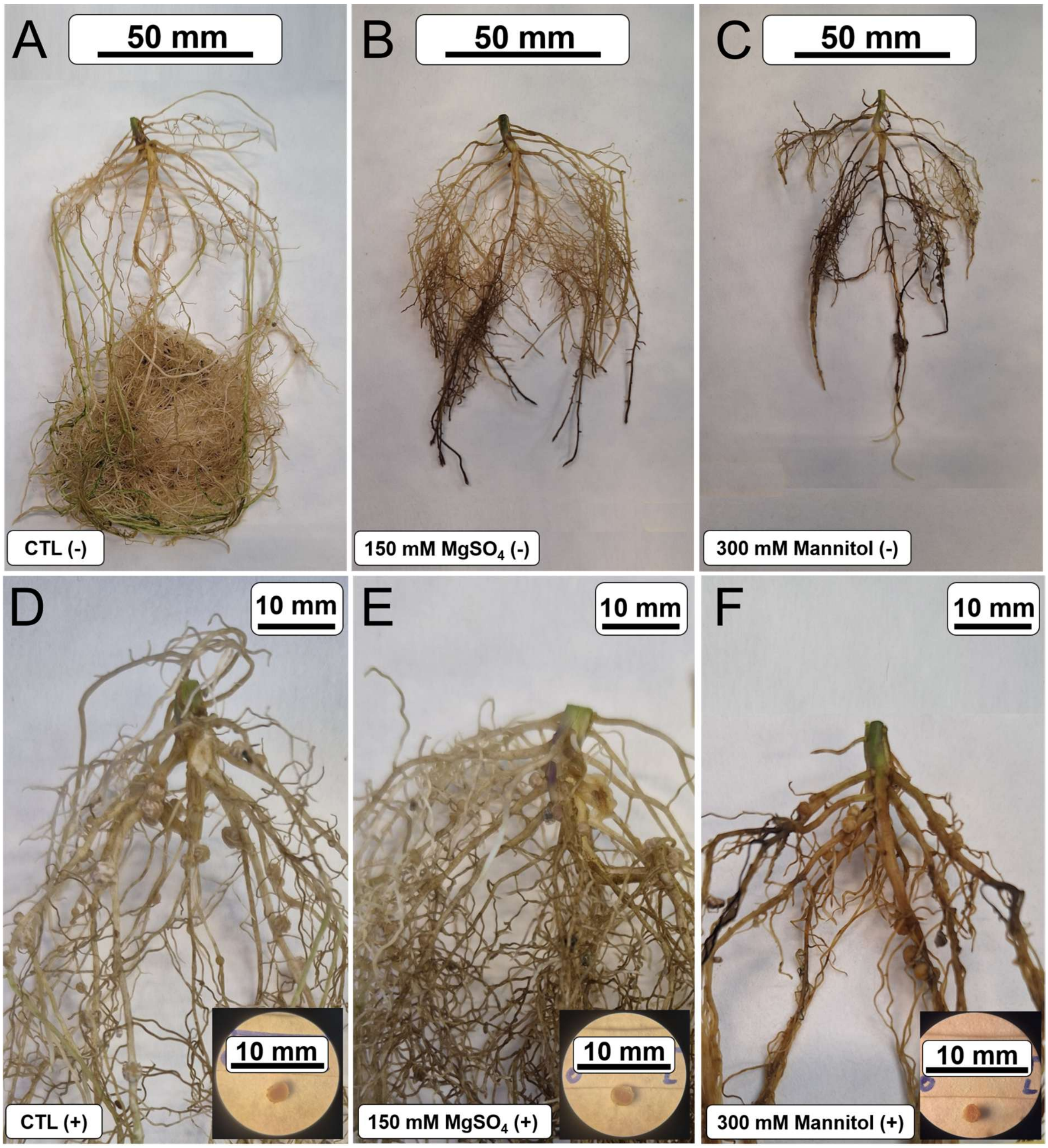


**Supplementary Figure 4** - **Effects of $MgSO_4$ and Mannitol on root system morphology of soybean plants, inoculated or not with *Bradyrhizobium japonicum*.** Representative root system of **(A)** non-inoculated control (CTL - ); **(B)** non-inoculated 150 mM $MgSO_4$; **(C)** non-inoculated 300 mM Mannitol; **(D)** inoculated control (CTL + ); **(E)** inoculated 150 mM $MgSO_4$; **(F)** inoculated 300 mM Mannitol.

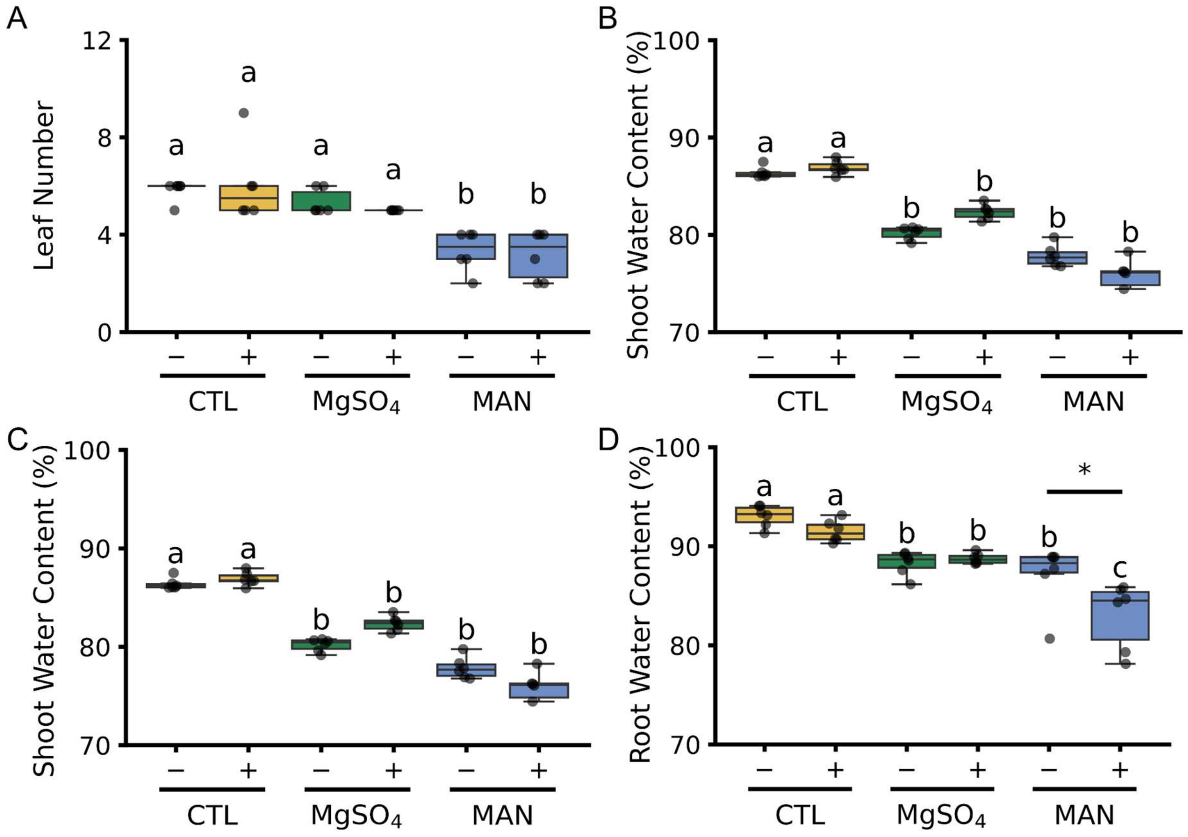


**Supplementary Figure 5 - Effects of $MgSO_4$ and Mannitol on growth parameters of soybean plants, inoculated or not with *Bradyrhizobium japonicum*.** **(A)** Leaf Number, **(B)** total water content, **(C)** shoot water content, **(D)** root water content. Plants were treated with water (yellow), 150 mM $MgSO_4$ (green) and 300 mM Mannitol (blue), with inoculation status indicated by (+) or (–) (n = 6). Lowercase letters denote significant differences among treatments (two-way ANOVA with Tukey's post hoc test, $p < 0.05$). At the same time, asterisks (*) indicate significant differences between inoculated and non-inoculated plants within the same substrate ($p < 0.05$).

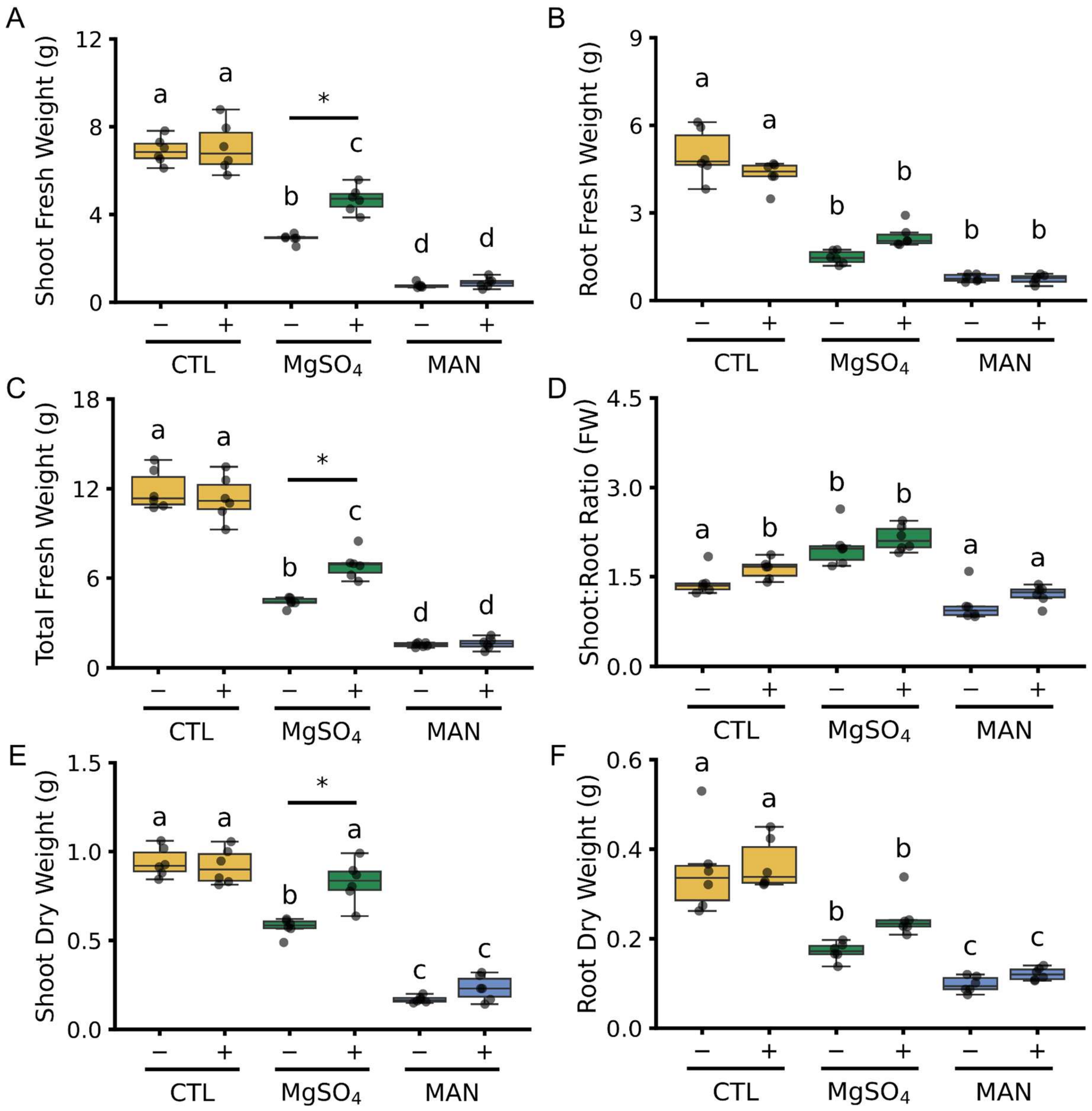


**Supplementary Figure 6 - Effects of $MgSO_4$ and Mannitol on growth parameters of soybean plants, inoculated or not with *Bradyrhizobium japonicum*.** **(A)** Shoot fresh weight, **(B)** root fresh weight, **(C)** total fresh weight, **(D)** shoot-to-root fresh weight ratio, **(E)** shoot dry weight, **(F)** root dry weight, **(G)** leaf number, and **(H)** total water content. Plants were treated with water (yellow), 150 mM $MgSO_4$ (green) and 300 mM Mannitol (blue), with inoculation status indicated by (+) or (–) (n = 6). Lowercase letters denote significant differences among treatments (two-way ANOVA with Tukey's post hoc test, $p < 0.05$). At the same time, asterisks (*) indicate significant differences between inoculated and non-inoculated plants within the same substrate ($p < 0.05$).

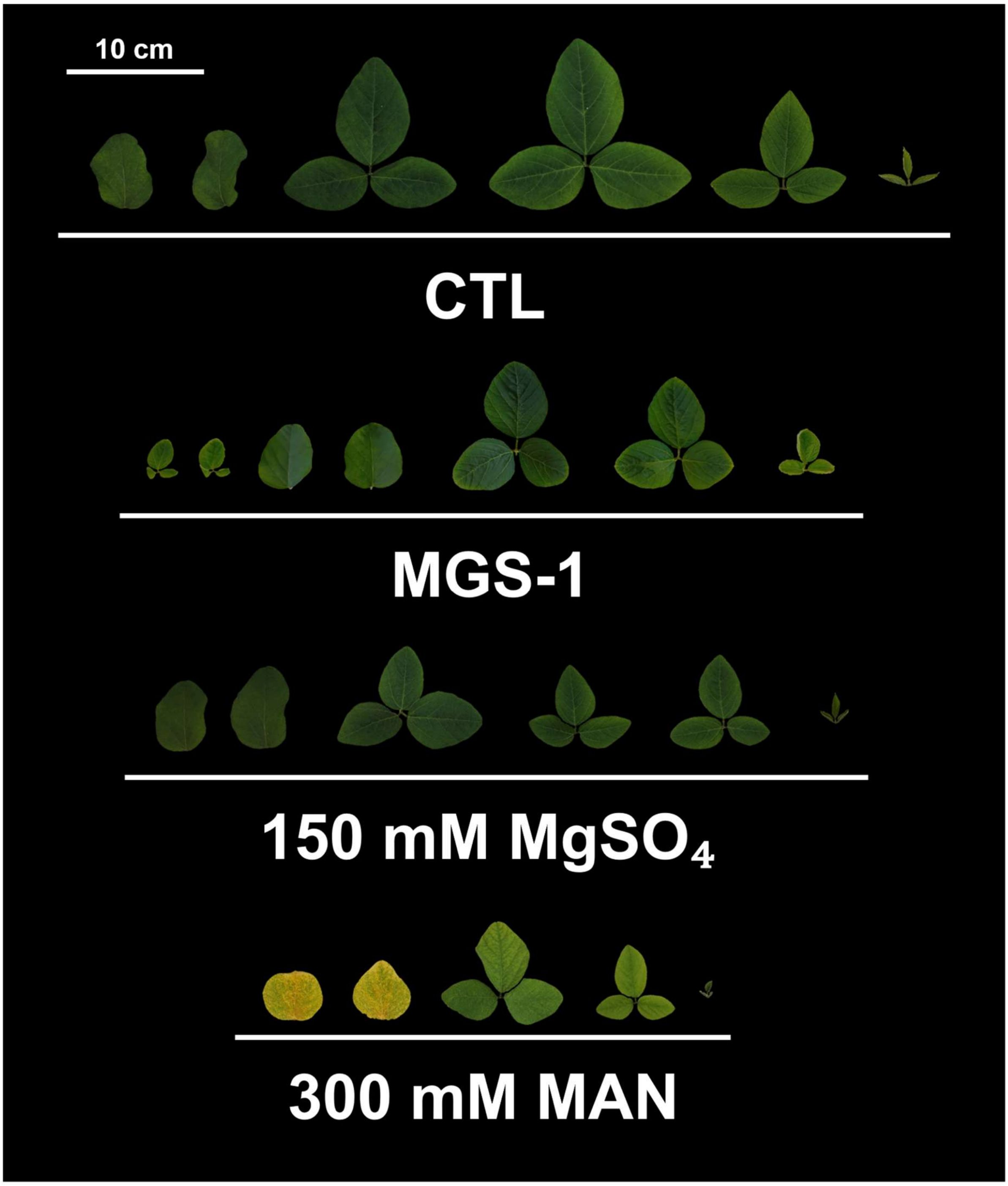


**Supplementary Figure 7 - Effects of MGS-1, $MgSO_4$, and Mannitol on leaf morphology of soybean plants.** Representative leaf morphology of soybean plants grown on sand without additions (control – CTL); MGS-1; sand supplemented with 150 mM $MgSO_4$; or sand supplemented with 300 mM Mannitol (from top to bottom, in this order), 28 days after imbibition (DAI). The horizontal white bar above CTL leaves represents 10 cm.

**Supplemental Table 1 – Results of two-way ANOVA examining the effects of MGS-1 and inoculation with Bradyrhizobium on plant growth parameters.** Degrees of freedom for each factor and the interaction are F(1, 8). For each dependent variable, the F-value, partial eta-squared ($\eta^2p$), and statistical significance are shown. * $p < 0.05$.

| | Substrate | | Inoculation | | Interaction (S*I) | |
|---|---|---|---|---|---|---|
| | F (1,8) | $\eta^2_p$ | F (1,8) | $\eta^2_p$ | F (1,8) | $\eta^2_p$ |
| **Height** | 37.68* | 0.65 | 0.06 | <0.01 | 5.11* | 0.20 |
| **Leaf number** | 58.78* | 0.75 | 10.10 | 0.34 | 0.87 | 0.04 |
| **Shoot Fresh Weight** | 55.46* | 0.87 | 1.53 | 0.16 | 1.85 | 0.19 |
| **Root Fresh Weight** | 56.55* | 0.88 | 0.68 | 0.08 | 0.14 | 0.02 |
| **Total Fresh Weight** | 74.75* | 0.90 | 0.06 | <0.01 | 0.34 | 0.04 |
| **S: R ratio (FW)** | 68.52* | 0.90 | 2.01 | 0.20 | 0.12 | 0.02 |
| **Shoot Dry Weight** | 28.34* | 0.78 | 0.16 | 0.02 | 0.17 | 0.02 |
| **Root Dry Weight** | 18.75* | 0.70 | 1.14 | 0.12 | 0.90 | 0.10 |
| **Total Dry Weight** | 44.84* | 0.85 | 0.48 | 0.06 | 0.34 | 0.04 |
| **S: R ratio (DW)** | 68.52* | 0.90 | 2.01 | 0.20 | 0.12 | 0.02 |
| **Shoot Water Content** | 91.12* | 0.92 | 3.96 | 0.33 | 16.36* | 0.67 |
| **Root Water Content** | 8.79* | 0.52 | 0.92 | 0.10 | 1.13 | 0.12 |
| **Total Water Content** | 0.04 | <0.01 | 1.23 | 0.13 | 3.30 | 0.29 |
| **S: R ratio (WC)** | 24.22* | 0.75 | 0.35 | 0.04 | 0.06 | <0.01 |
| **Shoot TSP** | 20.55* | 0.72 | 0.32 | 0.04 | 16.25* | 0.67 |
| **Root TSP** | 46.60* | 0.85 | 1.34 | 0.14 | 3.90 | 0.33 |
| **Total Chlorophyll** | 4.60 | 0.37 | 0.76 | 0.09 | 0.04 | <0.01 |
| **Total Carotenoids** | 0.55 | 0.06 | 0.69 | 0.08 | 0.07 | <0.01 |

**Supplemental Table 2 - Results of two-way ANOVA examining the effects of [$MgSO_4$] and inoculation with Bradyrhizobium on plant growth parameters.** Degrees of freedom for each factor and the interaction are F(3, 40). For each dependent variable, the F-value, partial eta-squared ($\eta^2p$), and statistical significance are shown. * $p < 0.05$.

| | [$MgSO_4$] | | Inoculation | | Interaction (Mg*I) | |
|---|---|---|---|---|---|---|
| | F (3,40) | $\eta^2_p$ | F (1,40) | $\eta^2_p$ | F (3,40) | $\eta^2_p$ |
| **Nodule number** | 2.46* | 0.16 | 156.53* | 0.80 | 2.46 | 0.16 |
| **Height** | 35.77* | 0.73 | 5.65* | 0.12 | 1.05 | 0.07 |
| **Leaf number** | 18.87* | 0.59 | 0.95 | 0.02 | 3.23* | 0.19 |
| **Shoot Fresh Weight** | 34.31* | 0.72 | 7.32* | 0.15 | 2.82 | 0.17 |
| **Root Fresh Weight** | 35.11* | 0.72 | 0.50 | 0.01 | 2.25 | 0.14 |
| **Total Fresh Weight** | 40.23* | 0.75 | 2.83 | 0.07 | 2.70 | 0.17 |
| **S: R ratio (FW)** | 32.55* | 0.71 | 1.99 | 0.05 | 1.87 | 0.12 |
| **Shoot Dry Weight** | 9.34* | 0.41 | 0.77 | 0.02 | 3.64* | 0.21 |
| **Root Dry Weight** | 12.63* | 0.49 | <0.01 | <0.01 | 3.28* | 0.20 |
| **Total Dry Weight** | 14.26* | 0.52 | 0.16 | <0.01 | 3.35* | 0.20 |
| **S: R ratio (DW)** | 77.25* | 0.85 | 15.81* | 0.28 | 8.42* | 0.39 |
| **Shoot Water Content** | 27.36* | 0.67 | 18.36* | 0.31 | 0.71 | 0.05 |
| **Root Water Content** | 2.67 | 0.17 | 5.84* | 0.13 | 5.55 | <0.01 |
| **Total Water Content** | 3.33* | 0.20 | 9.06* | 0.18 | 4.12 | 0.24 |
| **S: R ratio (WC)** | 12.69* | 0.49 | 0.20 | <0.01 | 4.56* | 0.25 |
| **Shoot TSP** | 3.75* | 0.22 | 9.58* | 0.19 | 3.16* | 0.19 |
| **Root TSP** | 19.58* | 0.59 | 11.90* | 0.23 | 1.89 | 0.12 |
| **Total Chlorophyll** | 0.71 | 0.05 | 1.85 | 0.04 | 13.48* | 0.50 |
| **Total Carotenoids** | 0.50 | 0.04 | 13.66* | 0.25 | 15.96* | 0.54 |

**Supplemental Table 3 -Summary of ordinary least squares (OLS) regression models for the effects of $MgSO_4$ and Bradyrhizobium inoculation on different growth parameters.** $MgSO_4$ was considered a continuous predictor, and the inoculation was a categorical predictor. For each dependent variable, the model was fitted as: Dependent ~ $MgSO_4$ + C(Inoculum). The table presents the adjusted $R^2$ (Adj. $R^2$), overall model F-statistic with degrees of freedom (F (2,45)), and unstandardized regression coefficients (coef) for $MgSO_4$, Inoculum, and their interaction (where applicable). The interaction term ($MgSO_4$: C(Inoculum)) was only included in models where it was statistically significant; otherwise, it is marked as n.a. Significance levels: *p* < 0.05, **p* < 0.01.

| | Adj.$R^2$ | F (2,45) | ercept ef | culum coef | MgSO4 coef | Interaction coef |
|---|---|---|---|---|---|---|
| **Height** | 0.656 | 45.8** | 84.72** | -9.21* | -0.34** | n.a. |
| **Leaf Number** | 0.444 | 19.8** | 8.56** | 0.38 | -0.02** | n.a. |
| **Nodule Number** | 0.758 | 74.72** | 2.25 | 17.33** | -0.03* | n.a. |
| **SFW** | 0.653 | 45.28** | 12.68** | 1.63* | -0.05** | n.a. |
| **RFW** | 0.530 | 27.47** | 11.53** | 0.58 | -0.07** | n.a. |
| **TFW** | 0.617 | 38.80** | 24.21** | 2.21 | -0.12** | n.a. |
| **S: R Ratio (FW)** | 0.489 | 23.51** | 1.19** | -0.22 | 0.01** | n.a. |
| **SDW** | 0.316 | 11.87** | 2.67** | 0.15 | -0.01** | n.a. |
| **RDW** | 0.277 | 10.00** | 2.43** | -0.03 | -0.02** | n.a. |
| **TDW** | 0.343 | 13.28** | 5.20** | 0.18 | -0.02** | n.a. |
| **S: R Ratio (DW)** | 0.704 | 38.30** | 1.49** | -0.70 | 0.02** | 0.02** |
| **SWC** | 0.651 | 44.78** | 78.70** | 3.09** | -0.06** | n.a. |
| **RWC** | 0.266 | 6.69** | 82.05** | -3.58 | -0.01 | 0.011* |
| **TWC** | 0.329 | 8.70** | 80.74** | -0.72 | -0.06** | 0.05** |
| **S: R Ratio (WC)** | 0.343 | 13.26** | 1.01** | -0.01 | <0.01* | n.a. |
| **Shoot TSP** | 0.263 | 9.38** | 69.76** | 12.53** | -0.12** | n.a. |
| **Root TSP** | 0.485 | 23.16** | 11.17** | 10.05** | 0.18** | n.a. |
| **Total Chlorophyll** | 0.465 | 14.55** | 1.16** | 0.42** | <0.01** | 0.01** |
| **Total Carotenoids** | 0.573 | 22.03** | 0.20** | 0.06** | <0.01** | <0.01** |

* p < 0.05, ** p < 0.01

**Supplemental Table 4 - Results of two-way ANOVA examining the effects of [$MgSO_4$], Mannitol and inoculation with Bradyrhizobium on plant growth parameters.** Degrees of freedom for each factor and the interaction are F(2, 30). For each dependent variable, the F-value, partial eta-squared ($\eta^2 p$), and statistical significance are shown. * $p < 0.05$.

| | **Treatment** | | **Inoculation** | | **Interaction (T*I)** | |
|---|---|---|---|---|---|---|
| | **F (2,30)** | $\eta^2_p$ | **F (1,30)** | $\eta^2_p$ | **F (2,40)** | $\eta^2_p$ |
| **Nodule number** | 6.80* | 0.31 | 158.98* | 0.84 | 6.80* | 0.31 |
| **Height** | 162.08* | 0.92 | 1.52 | 0.05 | 13.69* | 0.48 |
| **Leaf number** | 30.49* | 0.67 | 0.15 | <0.01 | 0.26 | 0.02 |
| **Shoot Fresh Weight** | 325.12* | 0.95 | 11.91* | 0.28 | 7.79* | 0.34 |
| **Root Fresh Weight** | 245.85* | 0.94 | <0.01 | <0.01 | 7.51* | 0.33 |
| **Total Fresh Weight** | 359.76* | 0.96 | 4.88* | 0.14 | 9.08* | 0.38 |
| **: R ratio (FW)** | 48.85* | 0.77 | 5.14* | 0.15 | 0.09 | <0.01 |
| **Shoot Dry Weight** | 257.97* | 0.95 | 13.20* | 0.31 | 9.27* | 0.38 |
| **Root Dry Weight** | 72.18* | 0.83 | 4.98* | 0.14 | 1.23 | 0.08 |
| **Total Dry Weight** | 214.79* | 0.93 | 12.15* | 0.29 | 6.66* | 0.31 |
| **S: R ratio (DW)** | 45.65* | 0.75 | 0.03 | <0.01 | 0.79 | 0.05 |
| **Shoot Water Content** | 39.28* | 0.72 | 0.36 | 0.01 | 3.92* | 0.21 |
| **Root Water Content** | 36.52* | 0.71 | 6.09* | 0.17 | 3.61* | 0.19 |
| **Total Water Content** | 114.56* | 0.88 | 7.20* | 0.19 | 15.03* | 0.50 |
| **S: R ratio (WC)** | 2.96 | 0.16 | 0.41 | 0.01 | 0.38 | 0.02 |

**Supplemental Table 5 – pH values of the substrates used before and after the experiments.** We measured the pH values of the substrates used according to the procedures described in the Handbook on Seedling Evaluation by the International Seed Testing Association (2008). We mixed 10 mL of each substrate with five volumes of sterile distilled water. Each substrate mixture was stirred continuously for 5 min, then allowed to stand for 2 h. Afterwards, each mixture was stirred, and the pH was measured using a pH meter (Sartorius PB-11, DWS Inc). To measure the pH of the substrates after the experiments, the substrates were dried in an oven at 60 °C. After drying, the substrates were maintained at room temperature, and pH values were measured as described above.

| Substrate | pH |
|---|---|
| Sand (before experiment) | 4.13 |
| Sand (after experiment) | 6.27 |
| Sand + *Bradyrhizobium* (after experiment) | 6.31 |
| MGS-1 (before experiment) | 7.83 |
| MGS-1 (after experiment) | 7.75 |
| MGS-1 + *Bradyrhizobium* (after experiment) | 7.86 |
